# Comparison of motility of H. pylori in broth and mucin reveals the interplay of effect of acid on bacterium and the rheology of the medium it swims in

**DOI:** 10.1101/2020.04.15.042622

**Authors:** C. Su, K. Bieniek, W. Liao, M. A. Constantino, S. M. Decker, B. S. Turner, R. Bansil

## Abstract

To colonize on the gastric epithelium *Helicobacter pylori* bacteria have to swim across a gradient of pH from 2-7 in the mucus layer. Previous studies of *H. pylori* motility have shown that at pH below 4 do not swim in porcine gastric mucin (PGM) gels. To separately assess the influence of gelation of PGM and that of pH on motors and pH sensitive receptors of *H. pylori*, we used phase contrast microscopy to compare the translational and rotational motion of *H. pylori* in PGM *versus* Brucella broth (BB10) at different pHs. We observed that decreasing pH leads to decreased fraction of motile swimmers with a decrease in the contribution of fast swimmers to the distributions of swimming speeds and length of trajectories. At all pH’s the bacteria swam faster with longer net displacement over the trajectory in BB10 as compared to PGM. While bacteria are stuck in PGM gels at low pH, they swim at low pH in broth, *albeit* with reduced speed. The body rotation rate and estimated cell body torque are weakly dependent on pH in BB10, whereas in PGM the torque increases with increasing viscosity and bacteria stuck in the low pH gel rotate faster than the motile bacteria. Our results show that *H. pylori* has optimal swimming under slightly acidic conditions, and exhibits mechanosensing when stuck in low pH mucin gels.

## INTRODUCTION

The human stomach presents one of the harshest environments due to the high acidity of its gastric juice secretion and various aspartate proteases and digestive enzymes which are crucial for metabolizing food and destroying microbes. To protect the stomach from its own acidic secretion and control the transport of food, microbes and other ingested products, the epithelial surface of the stomach is lined with a protective, continuous, viscoelastic layer of mucus varying from 100-400 μm in thickness. Across this mucus layer there exists a pH gradient maintained by the co-secretion of bicarbonate [1–3] pH near neutral close to the epithelial surface and highly acidic pH 2-4 on the luminal side during active acid secretion. The pH of the stomach measured at the luminal surface has been shown to range between 0.3 and 2.9 [4,5] with the resting median pH close to 1.74 [5] while the resting pH measured in the mucus layer has been shown to be close to 4 [4]. The mucus derives its viscoelastic properties from the glycoprotein mucin which has been shown to undergo a pH dependent sol to gel transition at pH 4 [6,7] forming a viscoelastic gel below pH 4. Exactly how the gelled mucin prevents the back diffusion of H+ ions is a subject of considerable debate and various processes such as H+ bindng to mucin, Donnan equilibrium, diffusion, viscous fingering have been invoked [8].

While the combination of gastric juice and the gastric mucus is quite effective in sterilizing and protecting the host from bacteria and infections, the gastrointestinal pathogen, *Helicobacter pylori* is known to breach this barrier and has adapted to this particularly challenging environment and colonized human stomachs [9]. *H. pylori* colonizes on the epithelial surface and even deep in the gastric glands, leading to the development of diseases such as gastric ulcers, gastritis, and even cancer. The role of stomach pH is one of the most important factors in controlling the colonization and pathogenic effects of *H. pylori* and many questions still remain poorly understood. Using their flagella-driven motility and further aided by chemotaxis mechanisms *H. pylori* navigates towards the neutral epithelial surface where it attaches using various adhesins and evokes an immune response and can cause inflammation of the surrounding tissue. Moreover, it is well known that *H. pylori* has anti-pH tactic chemoreceptor proteins that are sensitive to the environmental pH change, in particular, *TlpB* protein that are found in the inner membrane of *H. pylori* help it to move away from acidic pH [10]. In order to do so, the membrane-spanning *TlpB* proteins are activated by the protons in the environment, which triggers response of a cascade of chemotaxis proteins, such as CheY, CheW, and CheA [10]. The net response from the chemotaxis proteins contributes to the rotation direction and rate of the flagella motor, influencing the rate of reversal, persistence of track direction and speed of the bacteria. Furthermore, mucin itself a chemoattractant and is known to gel below pH 4 impeding the motility of the bacterium. Our group has previously reported that *H. pylori* shows no translational motion in porcine gastric mucin (PGM) solutions below pH 4 in the absence of urea, although rotation of their flagella could be observed in bacteria trapped in the mucin gel network [8,11]. The trapping of bacteria in the mucin gel is not surprising given that it has a structure of fiber bundles and pores at many length scales, ranging from around 200 nm as seen in AFM images of mucus [6] up to microns as seen in SEM images of mucus [8]. As is well known, *H. pylori* employs a urease secretion mechanism to hydrolyze urea which produces NH_3_ and CO_2_ [9]. In our previous work we show that the increase in pH above 4 due to urea hydrolysis leads to a gel to sol transition [11] enabling the bacteria to swim across the mucus layer. However, it is not clear from this previous study [11] how much of the observed pH dependent change in motility is due to the gelation of PGM and how much is due to the influence of pH on the flagellar motors, as the motors are driven by protons (H^+^), i.e. the proton motive force, PMF [12]. Unraveling the intertwined effect of pH on the bacterium and the medium through which it swims is clearly of importance to addressing how *H. pylori* breaches the pH and protective mucus barrier and to the development of strategies to control and treat the infection. Moreover, it presents an interesting case for the hydrodynamics of swimmers involving the coupling of pH effect on motors and pH sensing receptors along with a pH dependent change in the rheology of the medium transitioning from a sol to a gel. To the best of our knowledge the decoupling of these factors has not been addressed before.

The proton motive force (PMF) is related to the sum of the differences in pH, ΔpH, and membrane potential gradient ΔΨ across the inner membrane separating the cytoplasm from the periplasm (PMF = −61ΔpH + ΔΨ). For *H. pylori* the cytoplasmic pH which is regulated due to urease activity remains between 6.6 −7.6 for external pH between 5 and 8, and periplasmic pH is close to the external medium pH, as shown by direct measurements of both cytoplasmic and periplasmic pH using fluorescent dyes [13]. Both are relevant factors controlling pH dependence. Sachs *et al* measured the transmembrane potential for *H. pylori*, ΔΨ = PMF + 61ΔpH [14] finding a monotonic change in ΔΨ from −175 mV at pH 7.5 to −25 mV at pH 4 and becoming close to zero at pH 3.5. Combining these measurements leads to a PMF changing from around −130 mV at neutral pH [7,8] to −175 mV at pH 3. These data are also consistent with the observation that in the absence of urea *H. pylori* survives between a pH of 4.5 and 7.0 *in vitro*, and does not grow at a pH less than 3.5 without urea, i.e. it behaves as a neutrophile in standard buffers, not surviving extreme acidity or alkanity [15].

Previous studies of varying pH on motility in several bacteria such as *B. subtilis, E. coli* and *Salmonella* [16–19] have shown that motility is not affected by external pH but by the internal, i.e. cytoplasmic pH of the cell which can be modified by addition of a weak acid in the medium as these are known to permeate biological membranes. In addition to measuring the swimming speed the flagella rotation rate has been measured by tethering cells. For example in *E. coli* and *Salmonella* in the presence of a weak acid decreasing pH led to a decrease in swimming speeds over the pH range 7 to 5 and decrease in rotation rates [17] with the motor rotation stopping completely around pH 5. This was interpreted as an interference of the increase in intracellular H+ concentration with the protons released from the torque generating units. In contrast, *B.* subtilis swims well at pH 5.5 by maintaining a more-or-less constant PMF over the range of pH 5-7 arising mostly from ΔpH at pH 5 and ΔΨ at pH 7 [18]. Anti pH tactic behavior and enhancement of tumbling was observed in *Salmonella* in the presence of weak acids [16]. A study of torque-speed relationship showed that while high-speed rotation at low load was impaired in the presence of a weak acid [19] zero-speed torque was not affected. None of these effects were observed in the absence of weak acid. It is not clear whether these observations apply to *H. pylori* as it differs from *E. coli* and *B. subtilis* in having polytrichous, unipolar membrane-sheathed flagella. *H. pylori’s* motor has evolved to adapt to a highly viscous and somewhat acidic environment, using urease to regulate its cytoplasmic pH. It is capable of high torque generation having 18 stators per motor [20,21] as opposed to 11 in *E coli* and *Salmonella* and 8-11 in *B. subtilis.*

In this work our focus is to separately assess the influence of pH on loss of motility at low pH due to the viscoelasticity of the mucin gel and the influence of increased H+ concentration on the flagellar motors and pH sensing chemoreceptors of *H. pylori*. For this we compare the motility of *H. pylori* at different pHs in aqueous Brucella broth (BB10) *vs* PGM. While we have previously investigated motility and rheology at pH 4 and 6 [7,11] and there are some studies of the microrheology of PGM at neutral pH and pH 4 [22,23], there is no systematic study examining the pH dependence of the translational and rotational motion of *H. pylori* or the microrheology of mucin on length scales comparable to bacteria across the entire range of relevant pH’s. As expected, we observed that particle motion in broth is purely diffusive, in agreement with the bulk rheology studies reported previously [7], while that in PGM exhibits a strong pH-dependent sub-diffusion correlated with increasing heterogeneity at low pH. Live bacteria tracking shows that *H. pylori* swim faster in BB10 than PGM at all pH’s, and that motility is hindered at low pH in broth as well as in PGM. We find that decreasing pH primarily affects the faster swimmers and leads to a decrease in net displacement over the trajectory. These effects are more pronounced in BB10 as opposed to PGM presumably related to mucin binding to H+. In PGM at pH below 4 bacteria are immobilized due to the gelation of PGM, although they can still swim in BB10 at these low pHs. From the motion of individual bacteria at high magnification we observed that the body rotation speed *is higher* when they are immobilized in PGM at low pH as compared to the bacteria swimming at neutral pH, implying that perhaps bacteria are capable of sensing or coupling to the medium’s mechanical properties and rotate faster in the gel network. We also estimate torque at different pHs for each of the bacteria imaged at high magnification of 100X using Resistive Force Theory (RFT) following the methods of Magariyama *et al* [24] and found that in PGM bacteria have to generate a higher torque to overcome the increased viscosity, especially at low pH.

## RESULTS

### pH dependent microrheology of BB10 and PGM

To measure the microrheology of BB10 and PGM at various pH, we tracked several hundred to thousand trajectories from the Brownian motion of micron-size polystyrene latex particles suspended in the solutions using time-resolved fluorescence microscopy. The mean square displacement (MSD) versus time for each particle trajectory in BB10 and PGM at all pHs along with average <MSD> and relative error are shown in Supporting Information, Fig. S1. The viscosity, *η*, of each solution was computed as detailed in Su *et al* [25] using the time dependence of MSD for 2-dimensional diffusive motion of the particles,

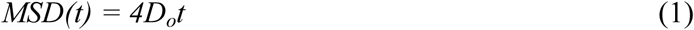

where *D*_*o*_ is the diffusion constant. The viscosity is inversely proportional to *D*_*o*_ via the Stokes-Einstein relation,

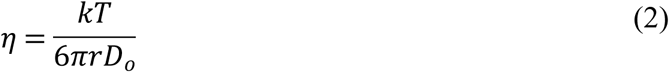

*k* is the Boltzmann constant, *T* is the temperature in Kelvin, and *r* is the radius of the particle. Fig 1**A** shows the <*MSD*> averaged over all particles in BB10 and PGM at various pH on a log-log plot, with water as reference. We found that the <*MSD*> and the viscosity of BB10 are similar to those of water, and not pH-dependent (Fig. 1**A, C**). In contrast in PGM, it is evident from the log-log plot of <*MSD*> that equation 1 is no longer valid and instead MSD is proportional to *t*^α^. By fitting the <*MSD*> *vs* time we obtained the exponent α < 1, indicating that the mobility of the micro-particles embedded in PGM is sub-diffusive, i.e. hindered at low pH as a result of the sol-gel transition that PGM undergoes as pH decreases below 4 [7,26]. As pH decreased from 6.7 to 3.7 in PGM, α deceased from 0.8 to 0.6, implying increasing sub-diffusivity of particles in PGM as it gels (Fig. 1**B**). In comparison to PGM, the microparticles in BB10 showed normal diffusion with α = 1 (Fig. 1**B**). Fig. 1**C** shows the viscosities of BB10 and 15mg/ml PGM calculated using Eqn. (2). In the case of PGM the effective viscosity was estimated by using only the data in the long-time regime where α ∼ 1. PGM is about 50 times more viscous than BB10 at pH 6, and the viscosity of PGM increased rapidly, by a factor of 2, as pH decreases from pH 6.1 to 3.7, whereas the viscosity of BB10 remains constant as pH decreases (Fig. 1**C**). Furthermore, in PGM the exponent α is time dependent, varying between 0.5 to 1, reflecting the frequency dependence of the moduli of the viscoelastic PGM [7,22,23]. The complex viscoelastic moduli were calculated and are reported in Constantino [27]. The viscosity estimated from the viscoelastic moduli agreed with that obtained from the MSD within 10%. Our results agree with the pH dependent trends observed in bulk rheology measurements using falling ball viscometry on 10 mg/ml PGM [28] showing a dramatic increase in viscosity as pH decreases. The viscosity of 10 mg/ml PGM at pH 4 is roughly 15 cP, as pH decreases from pH 4 to 2 the viscosity increases by a factor of 20 [28]. At low pH in PGM the elastic modulus approaches and becomes greater than the viscous modulus at low frequencies indicating gelation, although the moduli obtained from microrheology [8,23,25,27] are lower than those obtained from oscillatory shear bulk rheology [7,22] or falling ball microviscometry [28] showing that particles smaller than mesh size do not experience the bulk viscoelasticity. The micro- vs bulk methods also differ in time scales and shear rates probed. We also note that variation in viscosity values reported by different researchers is mostly due to differences in mucin purification processes, mucin composition and concentration and the sources from which the mucin was collected. However, one can clearly conclude that the overall viscosity increases with increasing mucin concentration and decreasing pH. We note that is important to obtain measurements of viscosity and motility from the same set of bacteria in the same batch of purified PGM to provide the most reliable comparison.

**Figure 1.**
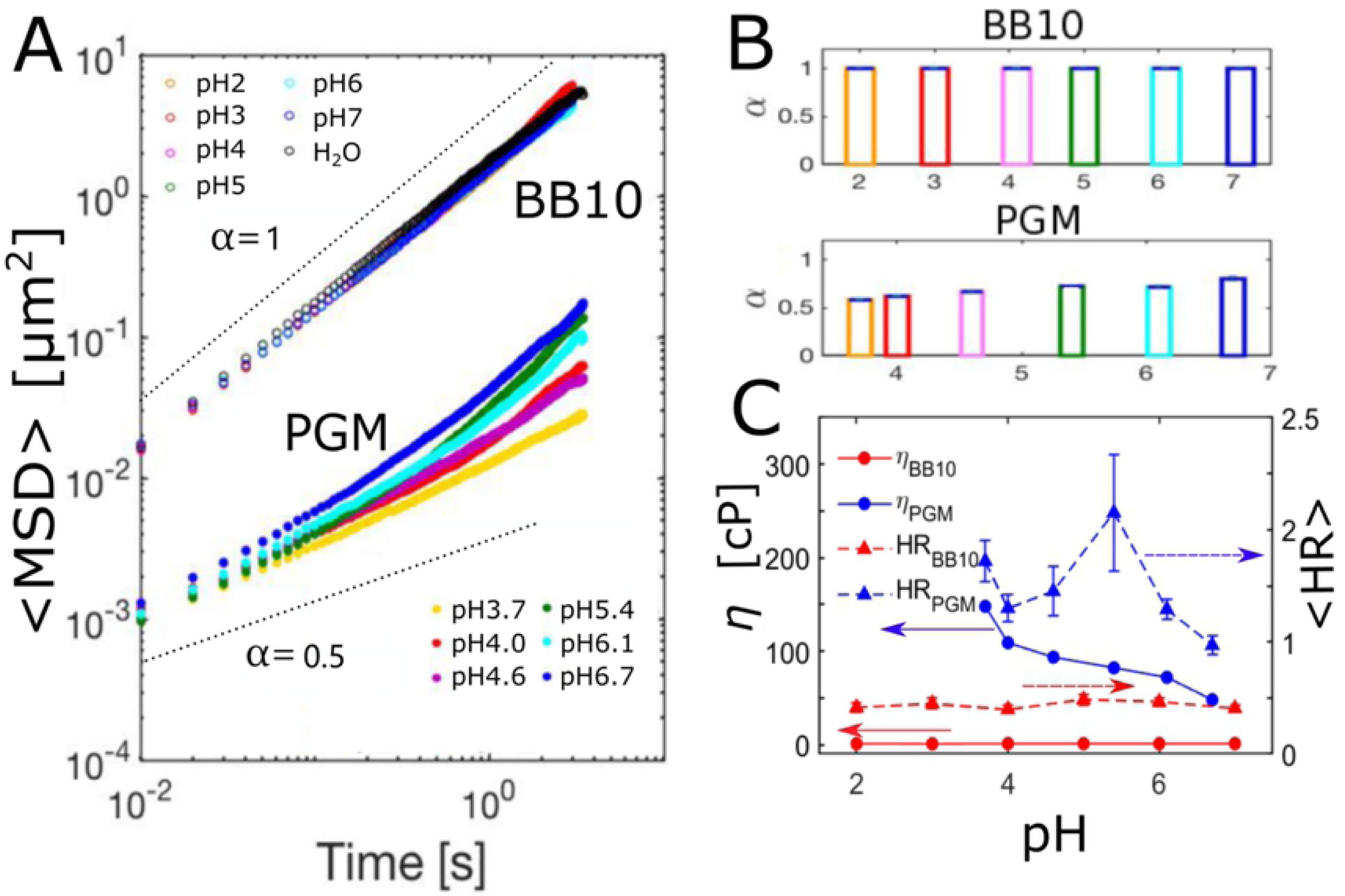
Microrheology of BB10 and PGM at various pHs. (**A**) Average mean-square displacement (<*MSD*>) of BB10 (open symbols), PGM (filled symbols) at pH 2 - 7 averaged over all particles. (**B**) Bar graphs of the exponent α *versus* pH calculated from <*MSD*> of all particles. Lines with slope α = 1 and 0.5 are drawn to guide the reader. (**C**) <*HR* > and viscosity *η* of BB10 and 15 mg/ml PGM calculated as discussed in the text. The arrows in (**C**) point towards the respective axes.

We evaluate the effect of gelation on mucin by computing the lag-time-dependent spatial heterogeneity (*HR*) of PGM at various pH following the methods described by Rich *et al* [30]. Briefly, *HR* is a dimensionless measure of the spatial heterogeneity in a non-homogeneous system, defined by the ensemble variance and the mean of *MSD*,

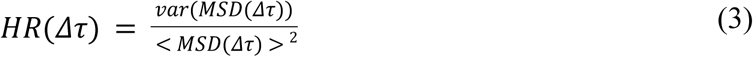

Here *Δτ* is the characteristic lag time. In particle tracking measurements due to particles constantly diffusing in and out of the focal plane, the ensemble averaging of *MSD* across all particles in a medium with spatially heterogeneous rheology tends to result in a statistical bias toward more motile particles, as they have higher probability of leaving and re-entering the field of view, producing segmented, shorter tracks. By calculating *HR*, each trajectory is weighted by a factor proportional to its duration in time. Fig. 1 C shows the time averaged <HR> along with its standard deviation as a function of pH in BB10 and PGM. As a reference from a theoretical calculation, in a bimodal system consisting of 50% water and 50% Newtonian fluid with viscosity ten times greater than water, *HR* ∼ 0.6 [30] at a characteristic lag time of *Δτ* = 0.1s, We found that *HR*_*BB10*_ ∼ 0.1 at all pHs measured, consistent with BB10 behaving like a watery Newtonian fluid, whereas at *Δτ* = 0.1s *HR*_*PGM*_ increases from ∼0.5 to nearly 1 as pH decreased from 6.7 to 3.7 indicative of increasing heterogeneity as pH decreases. The *HR* data in PGM shows a peak at pH 5, we suspect that this reflects a biphasic behavior of *HR*, with two characteristic trends corresponding to the gel phase at pH 4 and lower and the solution phase at pH 5 and above. From AFM measurements [6] we have shown that PGM fibers which are uniformly distributed at pH 6 begin to aggregate at pH 5, perhaps this leads to an increase in <HR> as particles encounter varying polymer concentration in different regions. In the gel phase, the particles are excluded from the regions with large concentration of fiber bundles and exhibit hindered motion confined in the pores of the gel. In this case the inhomogeneity will depend on the extent of crosslinking and degree of gel swelling/shrinking.

### pH dependence of swimming speed distributions of *H. pylori* in BB10 and PGM

To determine how the pH and the viscoelasticity of the environment influence the motility of *H. pylori*, we tracked bacteria from the *H. pylori* J99 strain swimming in BB10 and 15 mg/ml PGM with pH ranging from 2 to 6.3 using phase contrast microscopy recording time-lapse images at 40x magnification and 33 fps. We performed a detailed analysis of the movies to obtain the bacteria trajectories at each pH in BB10 and 15mg/ml PGM, following the methods described in Martinez *et al* [29]. Briefly, the PolyParticle Tracker program analyzes each trajectory to determine the instantaneous position as a function of time. Fig. 2 shows images of all recorded trajectories from entire movies at different pHs in BB10 (Fig. 2A) and in PGM (Fig. 2B). We note that the color coding of trajectories is not identified with the time (or the frame number in movies) where the trajectory began, although trajectories which cross each other were clearly from bacteria moving at different times. Different trajectories could arise from the same bacterium going in and out of the image plane; however, the program does not keep track of which bacterium gave which trajectory. We address this point in the last section by doing single bacteria tracking at high magnification. In the images of Fig. 2 we are displaying only the trajectories from swimming bacteria, those that were not motile were separately counted. From these images some differences between BB10 and PGM are clearly obvious. First comparing BB10 and PGM, we observe a larger fraction of linear trajectories in BB10 as compared to PGM. More trajectories in PGM exhibit a 2-dimensional random walk characteristic than in BB10.

**Figure 2:**
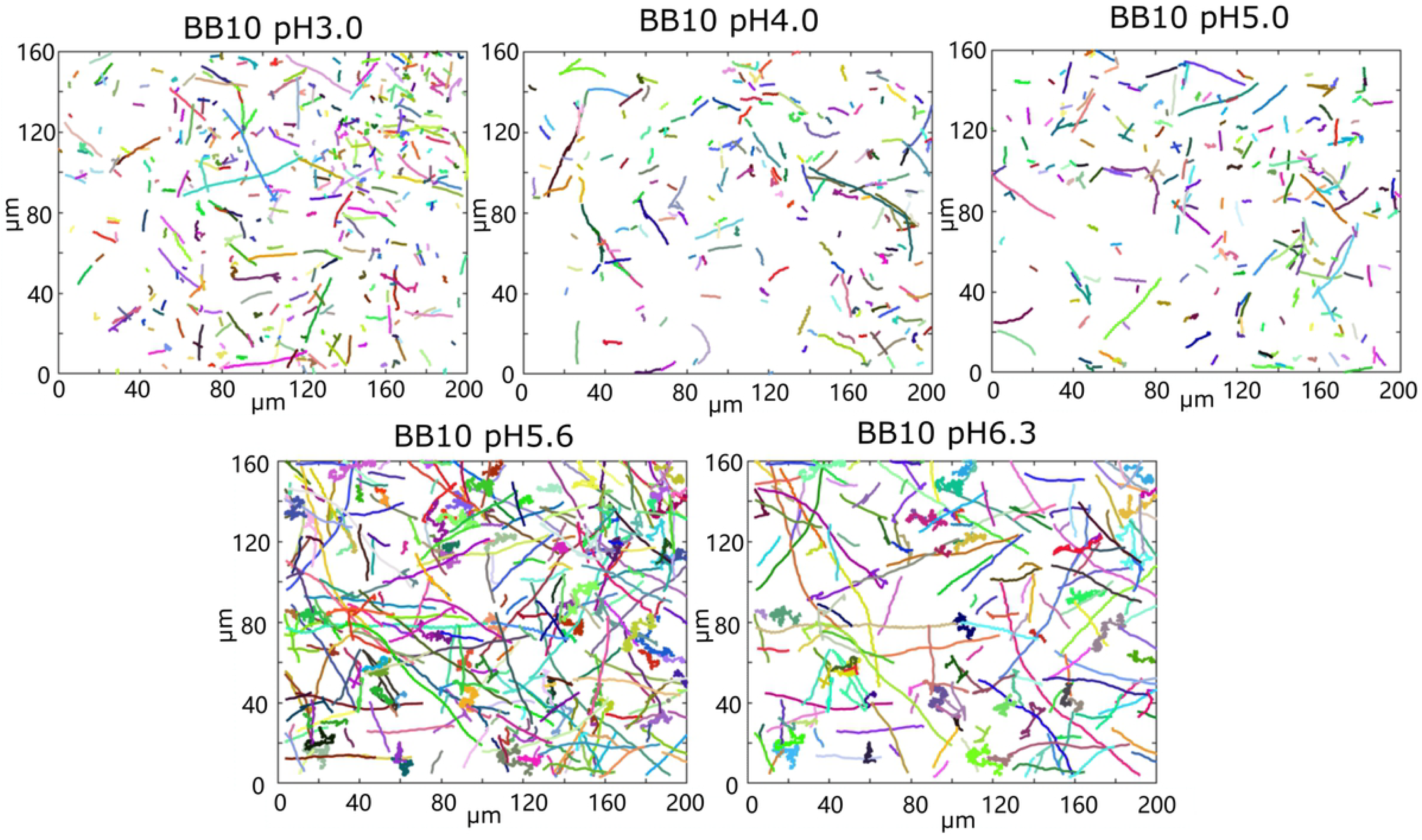
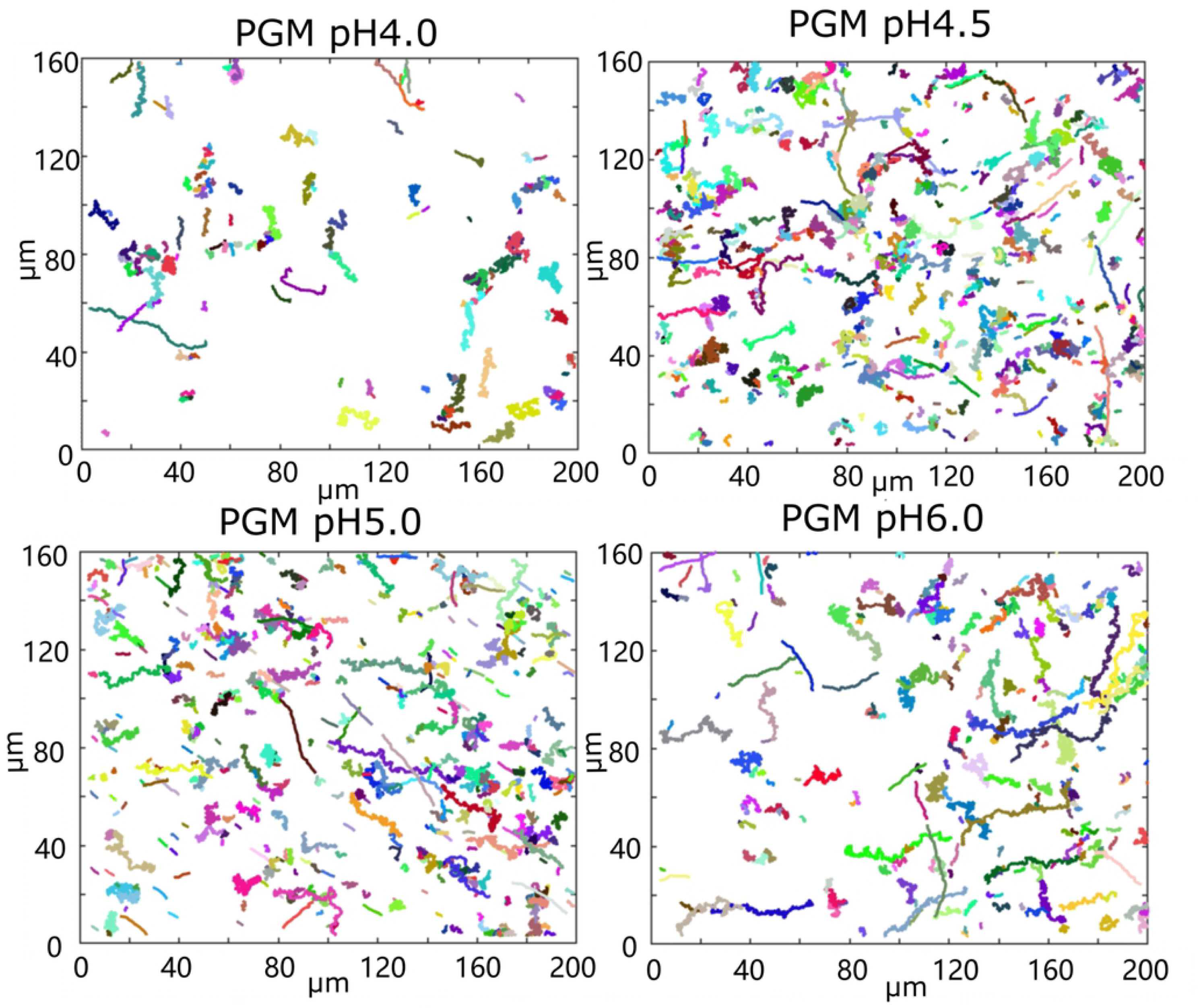
Images of trajectories in BB10 (A) and PGM (B) at different pHs. The trajectory images were created by stacking frames from 9s long video at 33fps to make a composite image.

Trajectories with turns and reversals can also be seen. The trajectories in PGM appear more helical than in BB10. We further discuss this helical feature in the last section using single bacteria imaging at 100 X magnification and fast frame rate to measure rotation of bacteria as they translate, similar to that reported by Constantino et al [31]. Secondly, regarding the effect of pH we note that in both BB10 and PGM there are considerably fewer trajectories at the low pH’s as compared to the images at pH > 5 in BB10 and pH > 4 in PGM. We found that in BB10 the bacteria swam over the entire range of pH 3 - 6.3, with a decline in the percentage of motile trajectories with decreasing pH, although some bacteria became immotile and coccoidal at pH 3. In contrast to this, in PGM the bacteria swam *only* at pH 4 and higher; there were very few swimmers (∼5 in total) in the pH 3.5 sample, and this data was not analyzed, while at pH 3 there were no swimmers; bacteria were stuck and observed to rotate in place at pH 3.5 and pH 3. The percent of motile bacteria was counted by looking at randomly selected frames in the movies. By this method we estimate that in BB10 there are about 40% motile bacteria at pH 3 and 4 and around 60% at pH 5 - 6, whereas in PGM there were only 12% motile bacteria at pH 4, and around 20-25% motile bacteria at pH 4.5, 5 and 6. The reference bacteria from the sample in BB10 were also examined at the same time as the PGM measurements were being conducted and these remained motile confirming that bacteria were viable and that the immotility was due to gelation of PGM not due to loss of motility in the sample.

From the trajectory we obtain the net displacement (*d*) between the first and last frame, as well as the maximum displacement (*d*_*max*_). For linear trajectories these two measures are the same, whereas for trajectories which look like directed random walks *d* < *d*_*max*_. Fig. 3 shows the histogram of the distribution of *d* for both BB10 and PGM at all pHs. The distribution of displacements extends to larger values for BB10 than PGM and this is reflected in the average <*d*> being about a factor of two larger in BB10 as compared to PGM. In both media, decreasing pH diminishes the occurrence of larger *d* values and this is reflected in <*d*> dropping by about a factor of 0.4 in BB10 between pH 6 and 5, and a more pronounced drop by a factor of 0.7 between pH 4.5 and 4 in PGM where no motile trajectories are found at pH 3. As another measure of the effect of pH on net displacement, we note that in BB10 at pH 5.6 and 6.3 over 40% of the trajectories are longer than 20 μm whereas this drops to 9-11 % at pH 5 and lower, and in PGM the corresponding numbers are smaller, about 21% at pH dropping to 7% at pH 5 and 4.5 and 3% at pH 4. The loss of motility with decreasing pH also manifests in fraction of trajectories which show active swimming *versus* passive Brownian motion. Active trajectories vary from over 95% at pH 5.6 and 6.3, and decrease to around 70% at pH 3 in BB10, whereas in PGM there are fewer active trajectories, ranging from about 25% at pH 6 to about 10% at pH 4. We attribute these differences between BB10 and PGM to the higher viscosity and ultimate gelation of PGM around pH 4.

**Figure 3.**
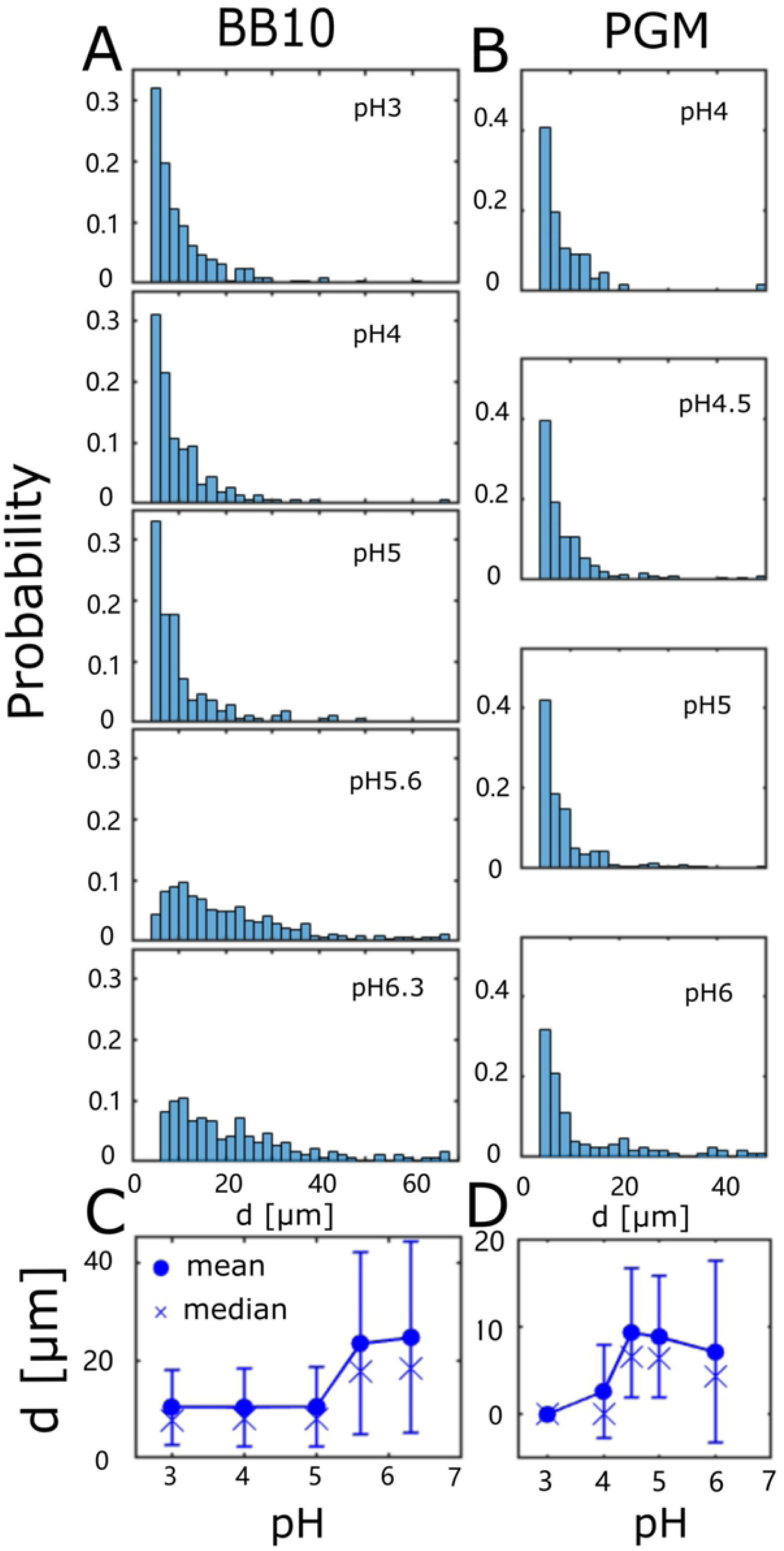
The distribution of trajectory displacements in BB10 (A) and PGM (B). Histogram of net displacement along a trajectory in BB10 (A) and PGM (B) at different pH’s as indicated, and the dependence of the average displacement (mean and median) on pH in BB10 (C) and PGM (D). Note that trajectories with *d* <4 um are not displayed as these correspond to bacteria which only display passive Brownian motion.

The trajectories were further analyzed by determining the instantaneous speed (*v*_*ins*_), defined as the displacement per unit time between two consecutive time frames, and the direction in which the bacterium was traveling by the change in angle of the displacement vector. Reorientation events were identified when bacteria changed the direction in which they were traveling, and when the reorientation angle (*θ*_*re*_) exceeds 140° we considered it as a reversal. The reversal frequency defined as the number of reversal events in a trajectory divided by the time duration of a trajectory and % reversals defined as the number of angle changes larger than 140 ° relative to the total number of all angle changes were calculated. The run speed (*v*_*run*_), is defined as the average speed over a linear path between two reorientations or reversal events. The distribution of both *v*_*ins*_ and *v*_*run*_ was obtained from the analysis of large number of trajectories, typically greater than 200, and in some cases up to 600 trajectories were analyzed. The average mean, median and standard deviation (σ) of *v*_*ins*_ and *v*_*run*_, as well as the percentage of motile trajectories were determined at each pH in BB10 and PGM. As discussed in our earlier work, the standard deviation σ is a measure of the width of the distribution and reflects both the temporal variation in speed, as well as the variation in speed largely due to the polydispersity in number of flagella as well as in helical shape and size of the bacteria in a given sample [29].

Fig 4 shows the histograms of distributions of *v*_*ins*_ and *v*_*run*_ in BB10 (**A**) and PGM (**B**) at different pHs. Both the speed distributions are broad and appear to be clearly bimodal in BB10, and asymmetric at high speeds in PGM. The faster swimmers are most likely the bacteria with larger number of flagella, as shown in ref. *30* by comparing mutants with on average one more and one less flagellum; although shape variation between individual cells also contributes to variation in speed. In BB10 (Fig 4 **A**) we can clearly see two peaks in *v*_*ins*_ distribution: a peak from slow swimmers in the vicinity of 10-15 μm/s and that of fast swimmers around 30-40 μm/s, and a high speed tail in distribution of *v*_*run*_ at pH 5.6 and 6.3. At lower pHs the peak corresponding to the faster swimming shifts from around 40 μm/s at pH above 5 to less than 30 μm/s at pH 5 and lower. At pH 5 the speed distributions in BB10 are very broad because the slow and fast peaks overlap, although two peaks are still visible in the distributions. At pH 5.6 and 6.3 there are several bacteria swimming at very high speeds (> 60 μm/s). The speed distribution for the slower bacteria appear to be less influenced by pH. In PGM the speeds are lower than in BB10 at all pHs and there are very few swimmers with speeds exceeding 30 μm/s in PGM (the probability is small so they cannot be identified in the histogram). However, the distribution of speeds displays an asymmetric profile at the higher speeds, indicating that bimodality of speeds is present, although to a much smaller extent than in BB10. We have fit the *v*_*ins*_ data to two peaks and the results presented in Supporting Information Fig. S2 agree with the statements made above by visual inspection of the speed distribution histograms. We also calculated the mean instantaneous and run speeds, < *v*_*ins*_> and < *v*_*run*_> as well as the median speeds and the standard deviation σ for the entire distribution as shown in (Fig. 4 C) for BB10 and PGM (Fig. 4D), respectively. Variation in the overall average is influenced by the 2 peak character of the distributions and is therefore not as straight-forward to relate to changing pH as the loss of the faster swimmers. In BB10 the average speeds of these broad distributions appear slightly dependent on pH, whereas in PGM the speed goes to zero at pH 3, but otherwise is little influenced by pH in the solution phase. In considering the effect of pH in PGM we have to consider the highly negatively charged polyelectrolytic property of mucin which binds H+ and influences the ratio of free vs bound H+ [32] and the final pH of the solution, as confirmed by our observation that the pH of PGM solutions is higher than the pH of the buffer added (see Materials and Methods and reference [27]).

**Figure 4.**
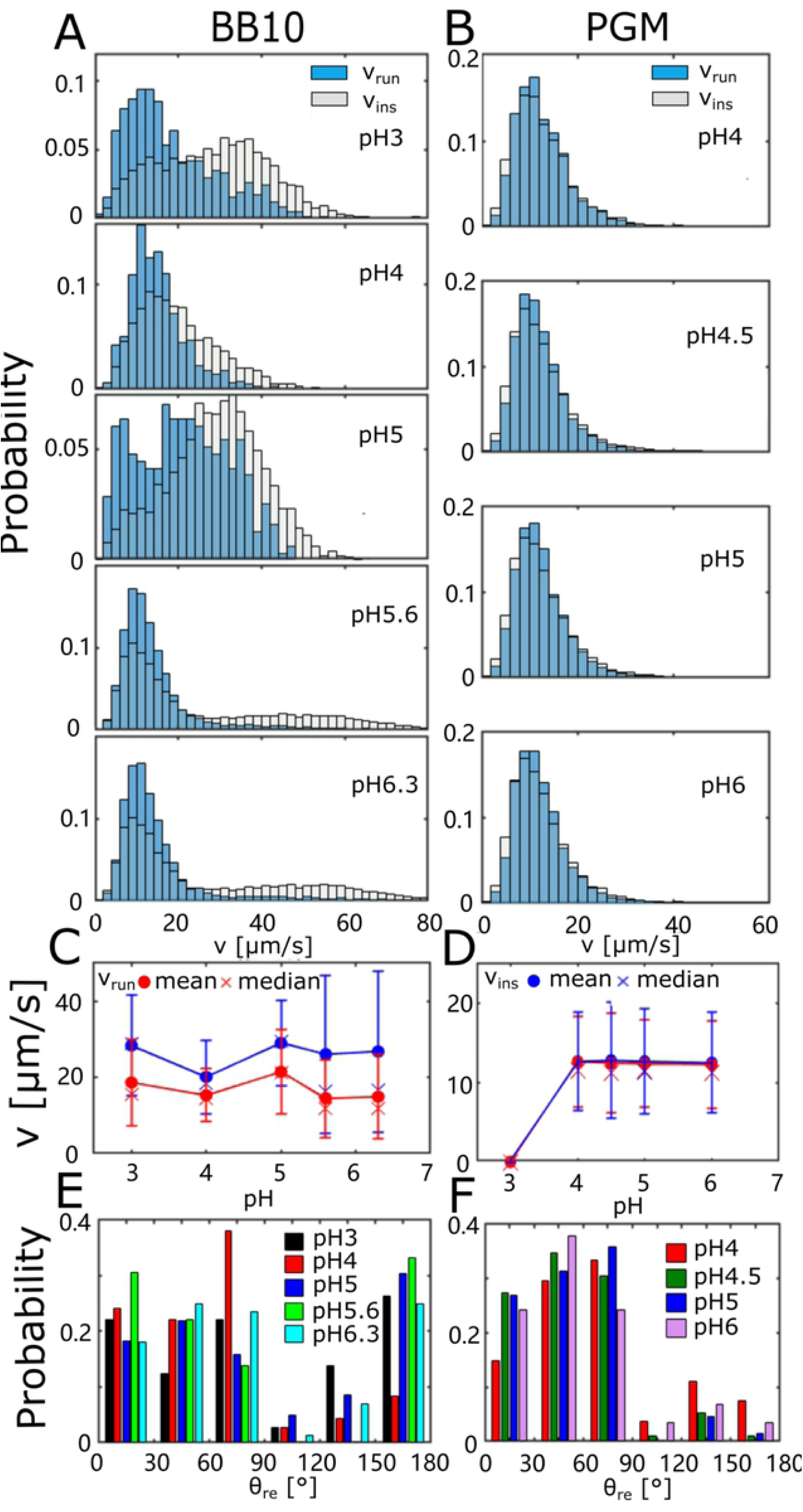
Distributions of swimming speeds and turn angles of *H. pylori* in BB10 and PGM at various pH. Histograms summarizing the instantaneous speed (*v*_*ins*_) (no color) and the run speed (*v*_*run*_) distributions (blue) in BB10 (**A**) and PGM (**B**) are shown at different pHs as indicated. When the two histograms overlap the outline of the *v*_*ins*_ histogram is visible over the blue histogram of *v*_*run*_. The average speeds (mean and median) for <*v*_*ins*_> and <*v*_*run*_> and standard deviation (σ) as a function of pH in BB10 and PGM are shown in **C** and **D**. The turn angle (*θ*_*re*_) distributions in BB10 (**E**) and in PGM (**F**) at different pHs as indicated.

The distribution of turn angles, *θ*_*re*_ at different pHs is shown in (Fig. 4 **E, F**) for BB10 and PGM, respectively. It clearly shows that there are fewer reversals in PGM at all pHs as compared to BB10, with a cumulative probability of reversals, estimated from turns between 140° −180°, ranging around 0.3 - 0.33 in BB10 (except at pH 4 which is low, around 0.13), and considerably smaller in PGM, ranging from about 0.03 in the solution phase (pH > 4) and increased to 0.07 in the gel phase at pH 3 (Fig. 4**F**). The reversal frequency decreases with decreasing pH in BB10 from 11/s to 3/s as pH decreases from 6.3 to 4, and is again high at pH 3 where the motor impairment occurs. On the other hand, in PGM the reversal frequency increases as pH decreases going from 1.5/s at high pH to 10/s at pH 4, reflecting the effects of increasing viscosity and gelation. Most of the reorientations in BB10 and PGM are less than 90° in both PGM and BB10, and this fraction of reorientations is higher in PGM than BB10 (cumulative probability for reorientation less than 90° is 0.6-0.8 in BB10 and 0.8-0.9 in PGM).

### The effect of pH on cell body rotation in BB10 and PGM

While there have been some previous results about effect of pH on swimming speeds, to the best of our knowledge there is no previous study about the effect of pH on *H. pylori’s* rotation. To measure the effect of pH on cell body rotation we imaged the motion of bacteria at a high magnification 100X and fast frame rate 100-200 fps as was previously done for *H. pylori* at neutral pH by Constantino *et al* [31]. Each frame from the trajectory of a single bacterium recorded as the bacterium rotates and translates was analyzed using the CellTool program [33] to obtain the cell body rotation. A few frames from one bacterium shown in Fig. 5**A** clearly shows that the bacterium projected shape as it is moving can be clearly imaged. As discussed in Constantino *et al* [31] the axis of the flagella and that of the cell body are not colinear, in other words the motion is like a precession. A typical contour and alignment axis of the bacterium obtained by CellTool is shown in Fig. 5**B**. This contour analysis was done for each frame of all the bacteria that could be imaged at different pHs in BB10 and in PGM. Fig 5**C** shows successive contours along the trajectory of the bacterium of Fig. 5**A** which shows the bacterium rotating while swimming forward as a pusher (A to B in the figure), then reversing to swim backwards (B to C) and reversing again at C to swim in the forward direction to the point D. The spacing of successive contours indicates that the bacteria is moving slower while it is in the reverse mode. In the movie (see Supporting Information Movie S1) we can see flagella sometimes which enables us to identify the pusher or puller (forward or reverse) motion. Fig 5**D** of alignment angle as a function of time shows clear oscillations as well as larger reorientations due to the trajectory changing direction between runs. From this data we can obtain the speed and frequency over each oscillation as well as the average speed *V* over the entire trajectory and the average rotation frequency *Ω*, also referred to as rotation rate, Because the images of cells are sufficiently clear at high magnification, when the bacterium image is fully in the image plane we can get good estimates of its length, thickness and helical pitch, and number of turns in the helix. We note that the speed measurement at this high magnification is from shorter trajectories than the ones reported in Fig. 2 from 40X imaging because the depth of focus is smaller and bacteria remain in focus for only short times. Moreover, some fast bacteria move out of focus so rapidly that they cannot be recorded. Due to these experimental limitations only a few bacteria could be tracked at high magnification and thus the swimming speed observed may not be statistically representative of the whole population of bacteria. In view of these limitations we focus on the body rotation rate from these high magnification measurements, which is quite constant over the short runs (see Fig. 5**D**). We also calculated the ratio *V/Ω* which is a measure of the distance travelled by the bacterium in one rotation. As discussed in the regularized Stokeslet modeling of the bacterium’s motion in Constantino *et al* the ratio *V/Ω* is independent of the flagella rotation rate [31]. The flagella rotation rate varies not only temporally but also from one bacterium to another as they have different number of flagella [29], and thus the ratio *V/Ω* is also independent of the number of flagella in each bacterium. Fig 5(**E, F**) show the bacteria cell body rotation rate (*Ω*) at different pHs in BB10 and PGM, respectively. By comparing bacteria in BB10 with those in PGM we note that while the rotation rate spans a similar overall range in PGM and BB10, the ratio *V/Ω* is about a factor of 5 smaller in PGM as compared to BB10 (Fig 5 **G, H**). This is consistent with the observation that due to the higher viscosity of PGM the bacteria swim slower in PGM than in BB10 (see the 40X measurements of Fig. 4). The rotation rates that we observe for J99 in pH 6 in BB10 and PGM are similar to those reported by Constantino *et al* [31] for the LSH100 strain which has similar average number of flagella as J99 [25,29,34]. A comparison of Fig. 5**E** and 5**F** shows that the rotation rate in BB10 is more or less unchanged by pH varying from pH 6 to 4 and then decreases as pH decreases to 3. The rotation rate in PGM exhibits a more complex behavior as below pH 4 the bacteria do not translate but still rotate [11]. In the pH range where bacteria can swim in PGM, the rotation rate decreases as pH decreases from 5.5 to 4.5, and then increases reaching a peak value at pH 3.5 and dropping slightly as pH drops to 3. The increase in body rotation frequency below pH 4 in PGM correlates with stuck bacteria suggesting an increased rotation speed of flagellar motors in a mechanosensing response to the increase in viscosity as pH approaches a sol to gel transition on decreasing pH below 4 (see microrheology data in Fig. 1). The rotation of stuck bacteria has previously been reported by Celli *et al* [11], although they were not able to compare rotation rates of stuck *vs* motile bacteria. We find that the values of the stuck bacteria rotation rate that we obtained are over one order of magnitude higher than the ∼3-10 Hz reported by Celli *et al* [11], which could just have been specific to the single bacterium they imaged, but also reflect the differences due to the different strain of *H. pylori* used in this study as compared to Celli *et al* [11] whose measurements were also at a slower frame rate (33 fps) and they only measured a couple of stuck bacteria. The movie reported by Celli *et al* [11] of the bacterium shows a large temporal variation in both body and flagella rotation rate, similar to our observations.

**Figure 5.**
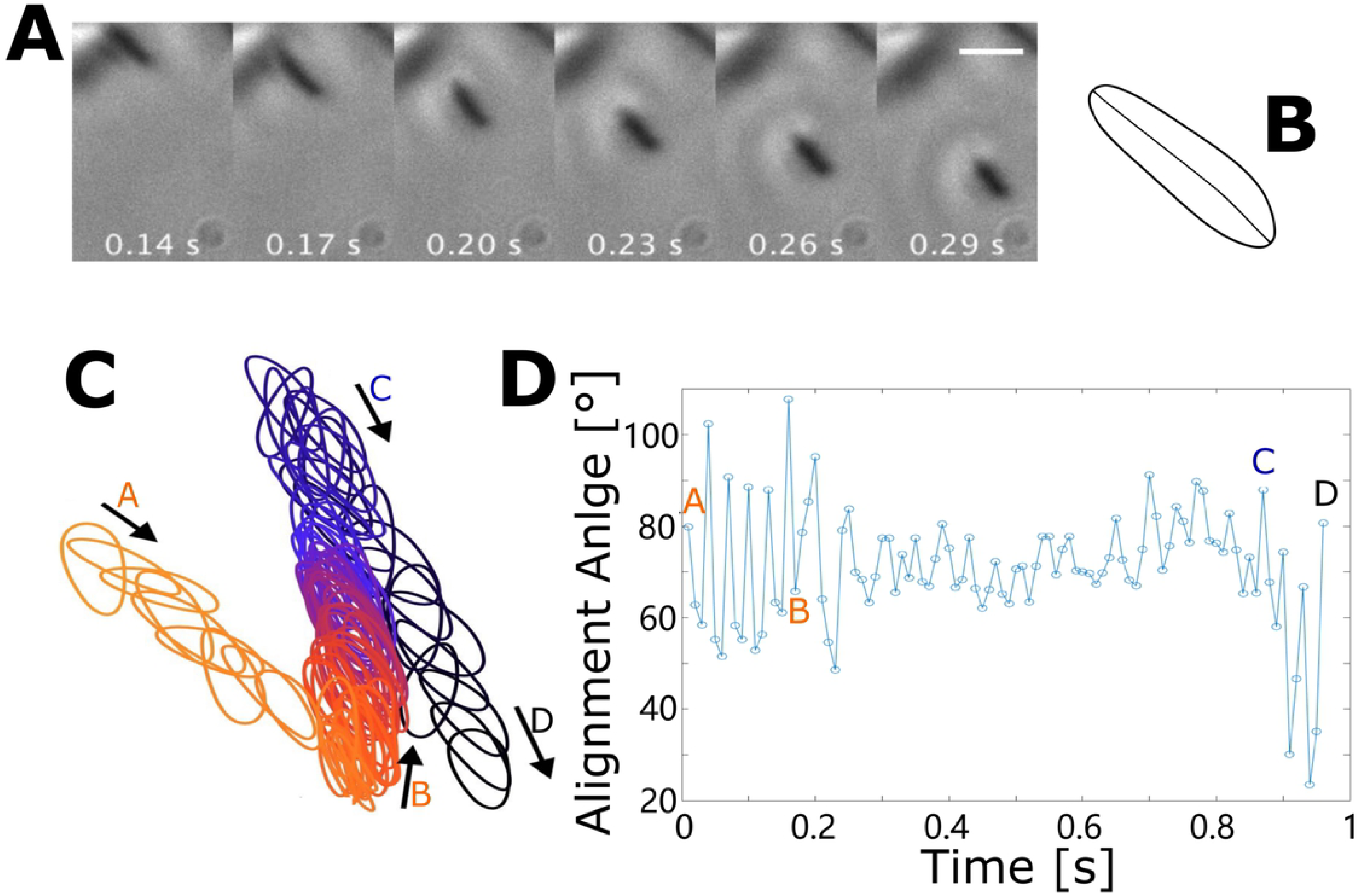
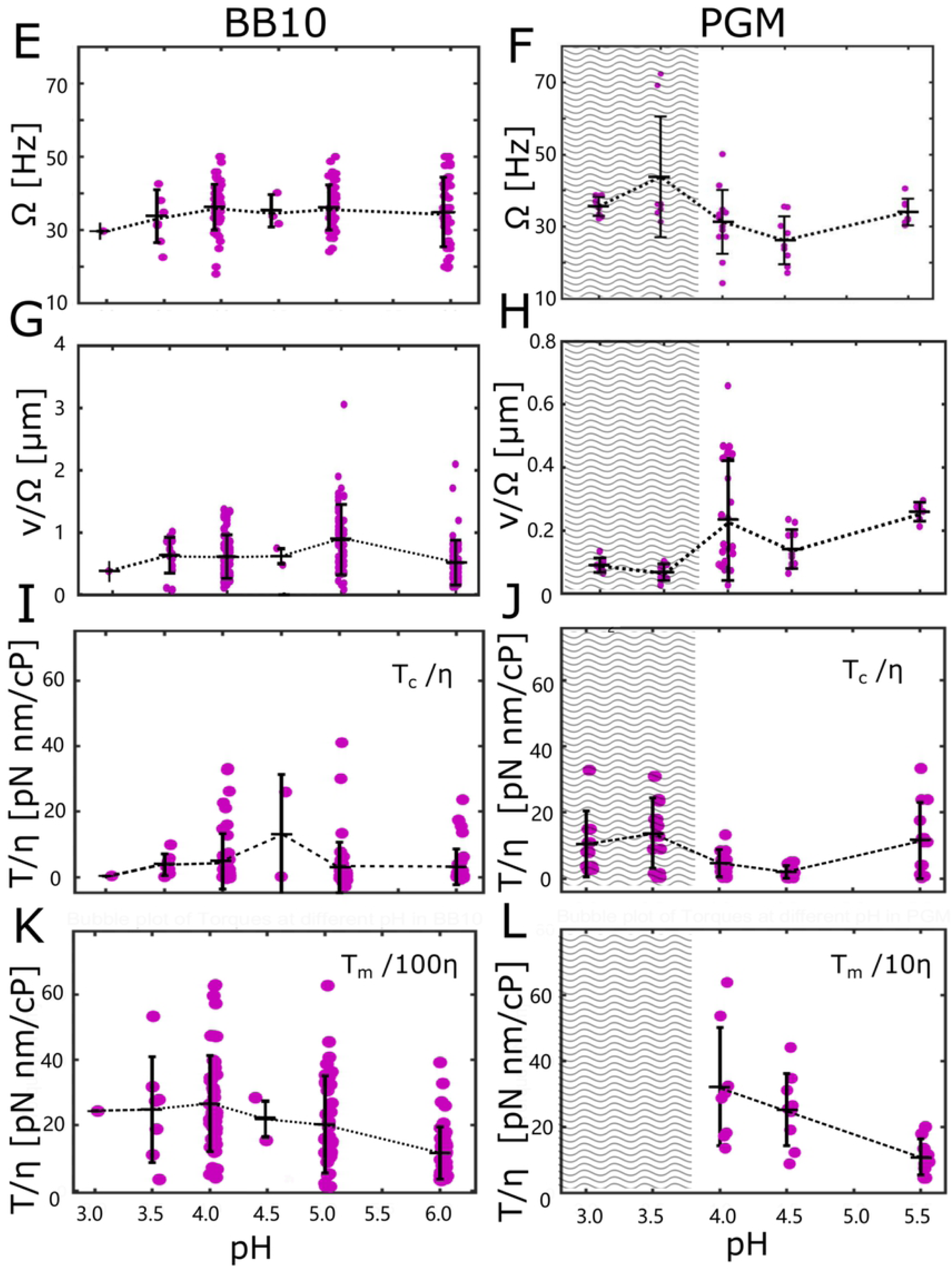
Rotation of cell body as *H. pylori* swims. (**A**) A series of frames showing the image of a single bacterium at 100X as it rotates while translating in BB10 at pH 4. (**B**) A typical contour and the center line obtained from CellTool. (**C**) Successive contours along the bacterium’s trajectory. The points A, B, C, D indicate reorientation events as described in the text. The frames shown in (**A**) are from the motion segment, A to B. (**D**) Alignment angle (with reference to an arbitrary direction) showing oscillations as the cell rotates. The movie for this data is provided in Supporting Information, (SI Movie S1). The bubble plots in **E, F, G, H, I, J** show cell body rotation rate (*Ω*) of the *H. pylori* bacteria in BB10 (**E**) and PGM (**F**), the ratio *V*/*Ω* in BB10 (**G**) and PGM (**H**), estimated cell body torque **T**_**c**_ in BB10 (**I**) and PGM (J), and motor torque **T**_**m**_ in BB10 (**K**) and PGM (**L**) calculated as described in the text for all of the bacteria imaged at pH 3 to 6. Note that **T**_**m**_ was not calculated for stuck bacteria for reasons discussed in the text. Cell body torque is plotted as T_c_ /η with torque in units of pN.nm and η in units of cP. The motor torque is plotted as T_m_ /(100 η) for BB10 and as T_m_ /(10 η) for PGM. The mean and standard deviation are indicated. The dashed lines are a guide to the eye. The gray shaded region represents the pH range over which PGM gels and bacteria did not translate but only rotated.

### Estimation of torque using Resistive Force Theory (RFT)

We estimated the torque for individual bacteria whose rotation rates and swimming speeds are presented in Fig. 5 **E - H** using RFT and a helical body as well as helical flagella bundle as was done in Martinez *et al* [29]. This is an improvement over the previous calculation in Celli *et al* [11] which was only done for stuck bacteria assuming an ellipsoidal body shape [11]. The RFT model we used is described in Magariyama *et al* [24]. Briefly, RFT decouples the flagella and cell body motion and thus forces and torques on an object are proportional to the local speed and angular rotation rate of that object, with the proportionality determined by the viscosity and the object’s geometry via hydrodynamic drag coefficients. The flagella and cell body satisfy force – torque balance and this enables calculation of the torques on both cell and flagella and the motor torque. We emphasize that RFT calculation is an estimate and has limitations both due to decoupling and assumptions such as implemented here with the flagella and body axis taken as colinear. More detailed studies could be done using Regularized Stokeslet model (RST) [24]. An additional caveat is that these RFT/RST models do not include viscoelasticity of the medium. Using the condition of zero net torque: *T*_*m*_ − *T*_*f*_ *= 0*, the motor torque *T*_*m*_, equal to the flagellar torque *T*_*f*_, can be estimated from bacterial translational speed *v*_*h*_ and shape factor *S*_*h.*_ For swimming bacteria, motor torque *T*_*m*_ was calculated from

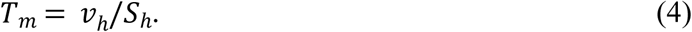

We used the measured speed for each bacterium imaged at 100X in PGM and BB10 at different pH and the shape factor *S*_*h*_ for a helical body and helical flagellum using the equations of Martinez *et al* [29] to calculate corresponding torque. (see the Methods section). The cell shape parameters of each bacterium were measured from the 100X images while the flagella parameters (not measured here) were taken to be the same as in Constantino *et al* [31]. The cell body torque *T*_*c*_ can be calculated from the measured swimming speed and angular velocity of cell body rotation ω _c_ = 2 π Ω where Ω is the measured rotation rate of the cell body.

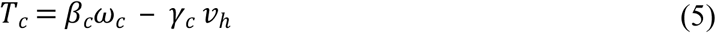

The hydrodynamic coefficients *β*_*c*_ and *γ*_*c*_ were obtained using the equations reported in Martinez *et al* [29] (see the Methods section for expressions for *β*_*c*_ and *γ*_*c*_). For stuck bacteria which rotate but do not translate *v*_*h*_ is zero and we obtain from Eq. (5),

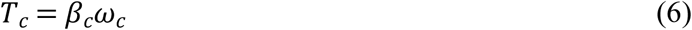

The hydrodynamic coefficients are all proportional to the viscosity *η* and thus torque is proportional to η. While η is close to 1 cP for BB10 and independent of pH (Fig. 1), the choice of what to use for η for PGM is complicated because mucin is a non-Newtonian, shear thinning fluid and at low pHs it is viscoelastic [7,27]. Furthermore, our previous studies of particle tracking microrheology of PGM in the presence of active, swimming *H. pylori* bacteria have shown that the probe particles exhibit enhanced diffusion due to the active swimmers [23,25]. The enhancement reflects the advection of particles as bacteria swim through the fluid, as well as the shear thinning that bacteria motion produces in PGM. These effects are more pronounced in the solution state at high pH and low concentration as compared to the gel state. Although advection of particles with swimming bacteria also occurs in BB10, there is no shear thinning and the overall enhancement in diffusion is not as high as in PGM. In our previous work we obtained a factor of 12.6 enhancement in diffusion of latex particles in the presence of J99 strain of *H. pylori* in PGM [25], implying an effective viscosity of PGM of 5 cP in the presence of bacteria swimming in PGM at pH 6. In view of these complications we plot the ratio *T*_*c*_ /*η* calculated using the equations above in Fig. 5 I and J for each of the bacteria imaged at 100X as a function of pH in BB10 and PGM, respectively. The calculation of *T*_*c*_ was done both for the stuck bacteria at low pH and the motile bacteria at higher pH. We find that the average ratio *T*_*c*_ /*η* in BB10 is comparable to that in PGM. However, the actual torque in PGM will be higher, scaling in proportion to the viscosity which increases as pH decreases (Fig. 1). If we use the actual viscosity of purified PGM [27] we get *T*_*c*_ to be about two orders of magnitude higher in PGM as compared to BB10. A more realistic estimate is obtained if we use the value of effective viscosity measured from particle tracking in PGM with active J99 *H. pylori* bacteria. In this case, torque *T*_*c*_ would be about 10-20 times higher in PGM than in BB10 at pH 6 and about 20-40 times higher in PGM at pH 4. For motor torque the estimates for BB10 shown in Fig. 5K increase from 1000 to 2000 pN. nm over the pH range 6 to 4 and remain more or less constant below that down to pH 3. On the other hand, in PGM using the viscosity of pure PGM from Fig. 1, motor torque would increase from a rather unrealistic value of 5000 pN. nm at pH 5.5 to 10,000 pN. nm at pH 4, whereas it would increase to more realistic values of 500-1000 pN.nm at pH 5.5 to about 1000-2000 pN.nm at pH 4, taking into account the reduction in effective viscosity of PGM due to shear thinning in PGM solution with active swimming bacteria.

In both media, the variation of torque among different bacteria imaged at the same pH is larger than the variation of the average with pH. The variation of cell torque *vs* pH reflects both the variation in *Ω* as well as the variation in size/shape of the cell and number of flagella for different bacteria imaged in the different measurements [29]. Torque is proportional to viscosity and hydrodynamic drag terms dependent on the size and shape of the bacterium. If all measurements were done with the same identical bacterium having identical flagella, then according to RFT, torque in BB10 would be independent of pH and that in PGM would increase with decreasing pH reflecting the increasing viscosity of PGM as pH decreases.

We used the flagella parameters from our previous Regularized Stokeslet modeling calculations [31], and estimated a motor torque about 100 times larger than cell body torque for motile bacteria in BB10 and 10 times larger in PGM (see Figs. 5**I** and **J**). For the case of stuck bacteria in PGM we cannot estimate the motor torque without additional assumptions because we could not measure the flagella rotation rate in our experiments. Furthermore, stuck bacteria which rotate but do not translate may have additional torques from the interactions between the bacterium and the medium which holds it in place.

### Different types of stuck bacteria

Fig 6 shows some images of bacteria stuck in different ways, obtained from the movies (S2 - S5) provided in Supporting Information. Along with a few images at different times as indicated, we also show the trajectory of the bacteria by plotting their positions over the entire movie as a single image (Figs. 6**E**-**H**). For e.g. one bacterium appears to be stuck at the flagella end (Fig. 6**A)**. Fig. 6**B** images show a pair of bacteria with one of them swimming in a small circle, while the other is in an orthogonal position. A bacterium at slightly higher pH 4.5 shows a small amount of translation with a few reorientations while it rotates (Fig. 6**C**). In contrast, the bacterium in BB10 at pH 4 clearly swims in a straight run (Fig. 6 **D**). Thus, it appears that bacteria can get stuck by having either their flagella or their body or both get entangled with the polymeric gel-like medium, or not be able to swim due to confinement in the pore of a gel, exhibiting passive, hindered Brownian motion.

**Figure 6.**
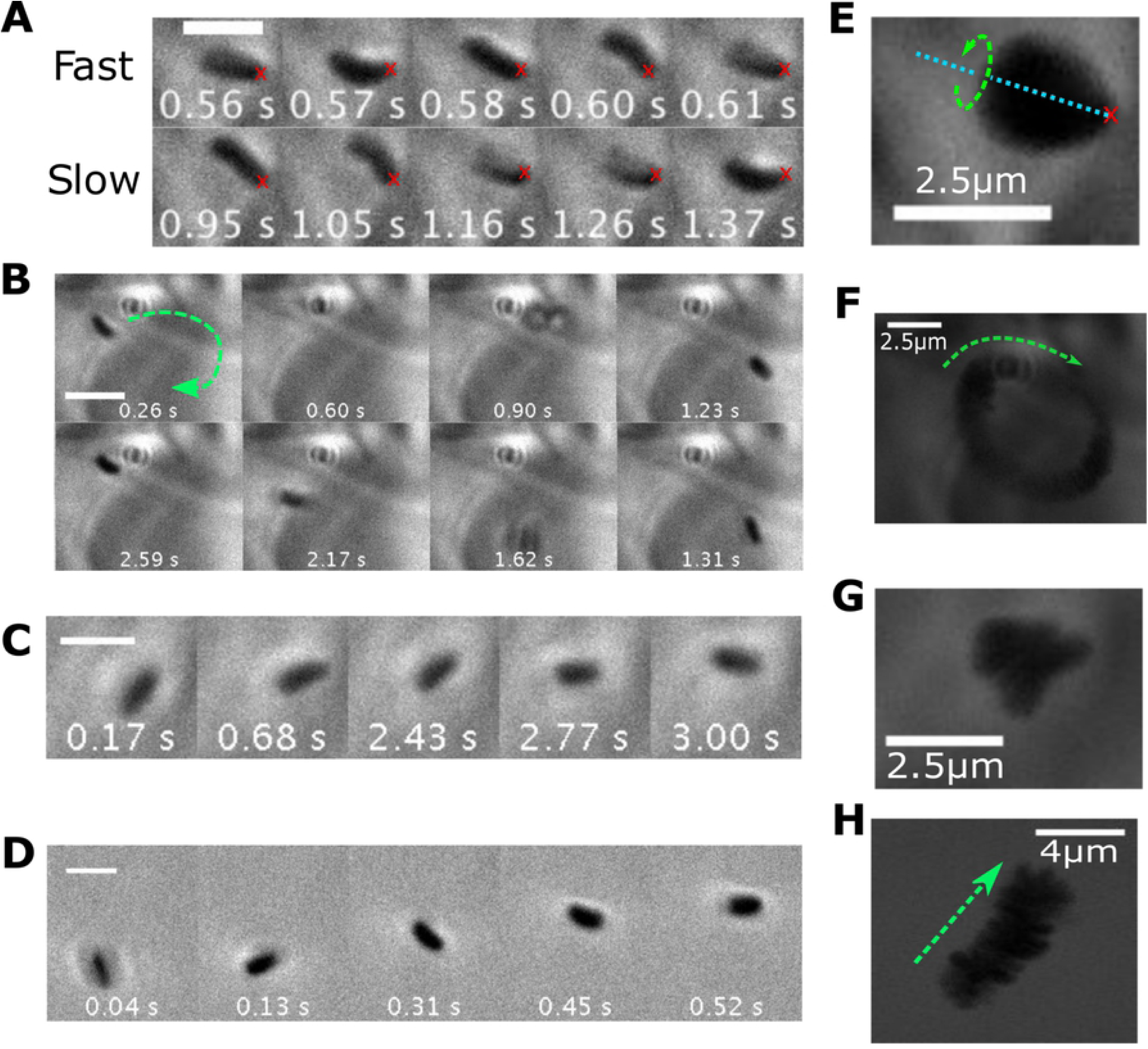
Phase contrast microscopic time-lapse montages and images of trajectories of various modes of *H. pylori* stuck in PGM at low pH. (**A**) Bacterium appears to rotate about a fixed point (indicated by the red x) with fast and slow body rotation rate over time. (**B**) Bacterium moves in a fixed circular trajectory perhaps reflecting proximity to a region of high PGM concentration (dark region) Time increases going clockwise through the images. (**C**) Bacterium at pH 4.5 in PGM showing rotations along with small, random translational displacements. (**D**) Bacterium in BB10 at pH 4 shows a clear translation while rotating. The scale bars in **A**-**D** indicate a length of 2.5 μm. The images in **E, F, G, H** are overlays of different frames of the movie onto a single image to show the trajectory corresponding to **A, B, C, D** respectively, with the green arrows indicating the direction of motion. The movies for this data are provided in Supporting Information, (Movies S2 – S5).

## DISCUSSION

By comparing the effects of varying pH and viscosity on the motility and body rotation of *H. pylori* in aqueous broth and PGM solutions, we find that while the flagella motors are impaired at low pH the effect of gelation of the medium plays a dominant role and totally immobilizes the bacteria. We observe that at all pH’s where the *H. pylori* bacteria are motile, they swim with about two times longer trajectories and about two times faster speeds in BB10 as compared to PGM. The decrease in distance traveled (net displacement) with decreasing pH in PGM correlates with increasing heterogeneity in PGM as pH decreases and viscosity increases. The swimming speed and trajectory displacement distribution extend to higher values in BB10 as compared to PGM and decreasing pH primarily diminishes the contribution of the longer and faster swimmers. The effect of varying pH on swimming speed in PGM was less pronounced than the effect in BB10. This may be related to PGM being a negatively charged polyelectrolyte so that H+ are trapped in the mucin gel [8]. Titration studies of PGM have shown that the fraction of free protons is reduced with decreasing pH [32], and the pH of the PGM solution is higher than that of the external buffer added.

The fraction of motile *H. pylori* bacteria declines with decreasing pH as observed for the other bacteria. Although there are fewer reversals in PGM as compared to broth, the reversal frequency increases at pH 4 in PGM, presumably reflecting the effect of gelation at low pH in PGM. Together with the speed distribution analysis, our results suggest that in BB10 we see a pH-dependence in *H. pylori* swimming speeds and reversal rates providing optimal swimming around a slightly acidic pH of 5. It is possible that *H. pylori* has adapted to transit through an acidic environment. There is other evidence which supports that *H. pylori* thrives in acidic environment, such as in presence of urea, an acidic environment was necessary for the survival of *H. pylori* [15] and the urease and metabolic activities of the bacteria increase at pH 4 and below [35]. *H. pylori* is also able to both survive and colonize deep in the parietal glands, where acid is secreted [36].

We found that the body rotation rate is only weakly dependent on pH in BB10 from pH 6 to 4 and then decreases at pH < 4, whereas in PGM it shows a peak at pH 3.5 correlated to viscosity increase as pH decreases. The decrease in the rotation rate at pH 3 in both PGM and BB10 reflects the impairment of flagella motors. At this low pH many bacteria were coccoidal in both BB10 and PGM indicating that under such extreme acidity bacteria are directly impacted by acid. In BB10 bacteria were observed to swim even at the lowest pH measured, whereas in PGM below pH 4 the bacteria are trapped in the gelled mucin and do not translate but only rotate in place, in agreement with previous reports [31]. In previous work from our group we have also observed stuck but rotating *H. pylori* bacteria in gelatin gels, but not in gelatin or methyl cellulose solutions [37]. The gelling of PGM or gelatin traps bacteria in liquid pockets or the bacteria cell body or flagella can be physically stuck in the gel network, even at a single point (as shown in Fig 6). The distribution of displacement shows that immotile bacteria also include bacteria doing passive, hindered Brownian motion in the pores of the gel. For stuck bacteria the interaction with the medium could lead to additional forces and torques which hold the bacteria in place and this might explain the increase in rotational frequency of the cell and loss of translational motion. We also observed that stuck bacteria rotate faster than motile ones, perhaps indicating a mechanosensing ability of *H. pylori* trapped in a gel network. Mechanosensing has been observed in recent studies of the flagellar motor with varying load and viscosity of the medium [38,39] and they suggest that the bacterial flagellar motor senses external load and mediates the strength of stator binding to the rest of the motor. The motor of *H. pylori*, like other ε-proteobacters such as *C. jejuni*, has adapted to high torque generation by having a large scaffold of proteinaceous, periplasmic rings to increase the radius of the contact lever point and to accommodate the large number of stators (17 for *C. jejuni* [21,40] and 18 for *H. pylori* [20,21], and by having high stator occupancy [21]. We can estimate the maximum torque for *H. pylori* using the same approach as Beeby et al [40] by assuming that torques of all stators are additive, and each stator complex exerts a force of 7.3 pN. For the lever arm we use 27 nm as the distance between the outer lobe of the C ring and the axis of rotation to obtain a maximum torque of 18 x 7.3pN x 27nm ∼ 3550 pN. nm per flagellum. As mentioned earlier, the unipolar bundle of the *H. pylori* J99 strain used here has 1-6 flagella with on average 3 flagella [34] similar to the LSH 100 strain used in our previous studies [29,31], implying that *H. pylori* could exert a torque of 1-6 times the maximal torque per flagellum, and that it has the capacity to vary its motor torque by 2 orders of magnitude by varying the number of active stators (from 1 to 6 x18 =108 stators). Furthermore, periplasmic pH which is close to external pH may directly impact motor function as the scaffolding rings are in the periplasmic space. The faster swimmers are more likely to have larger number of flagella [29] and as a consequence more stator units can be turned on. With decreasing external pH the proteins in the scaffold structure in the periplasmic space could be impacted and affect the stators. The direct influence of pH on these structures would be worth investigating.

The proton concentration in the surrounding medium has a direct effect on the swimming speed of *H. pylori* as it influences the transmembrane potential gradient and the difference in pH across the periplasmic membrane, as well as generating anti-pH tactic chemoreceptor responses. As mentioned earlier, previous studies on *E. coli, Salmonella* and *B. subtilis* [16–19] showed loss in motility and other changes at low pH’s in the presence of weak acids which can penetrate and alter the cytoplasmic pH, but no changes in the absence of weak acids. These results cannot be directly compared to our findings with *H. pylori* because we added HCl to change pH of the 0.1 M phosphate-succinate buffer composed of the weak phosphoric and succinic acids. Furthermore, in contrast to these other bacteria, *H. pylori* has evolved to live in an acidic niche [15] by utilizing urea hydrolysis and was able to swim even at low pH of 3 in BB10, although with reduced speed. As discussed in the Introduction, the work of Sachs et al and Wen et al [13,14] shows that the periplasmic pH of *H. pylori* is close to the external medium pH, while in the presence of urea the cytoplasmic pH is regulated due to urease activity and changes only slightly with external pH giving a Δ pH ranging from about 1 at pH 6 to about 3 at pH 3.5. Combining this with the measured transmembrane potential for *H. pylori*, ΔΨ changing from - 175 mV at pH 7.5 to −25 mV at pH 4 and becoming close to zero at pH 3.5 [14] leads to a PMF changing from around −130 mV at neutral pH [7,8] to −175 mV at pH 3. Our observation that when pH becomes too low (around pH 3) the flagellar motors do not function well is in agreement with the finding of Sachs *et al* [14], although our experiments were done without adding urea in the medium to prevent external pH from increasing [11]. The lack of added urea in broth or PGM medium could have impaired cytoplasmic pH regulation as the urease receptors on the external surface of the bacterium were probably not activated, although the internal ones probably were active [14,15,36].

In conclusion, our findings show that pH has a strong effect on the motility of *H. pylori*, reflecting the combined effects on flagellar motors and their immobility in the viscoelastic PGM gel as the bacteria get stuck and cannot translate even though they rotate faster as they mechanosense their environment. *H. pylori* is an ideal candidate for more detailed studies of pH dependence of torque-speed relationships as it provides an inherent coupling of the properties of the medium (pH dependent viscosity, gelation, specific adhesin binding and polyelectrolytic interactions) to effects of pH on motors and receptors, and the bacterium in turn influences the properties of the medium. Further investigation to reveal the correlation between the transmembrane proton gradient, pH effects on stators and scaffolding complex and activation of *TlpB, ChePep* and other flagellar proteins could explain why faster swimming bacteria at lower pH travel in straighter trajectories as shown in our results, and why lowering pH predominantly affects the faster swimming bacteria.

Our observations lend further insight into the mechanism by which *H. pylori* is able to breach the gastric mucus barrier and colonize on the gastric epithelium causing diseases such as gastritis, gastric ulcers and even leading to gastric cancer. The work has potential relevance to regulation of acid in the treatment of *H. pylori* infection. Preliminary measurements of chemotactic motility under pH gradients are reported in [20,41] and provide a more realistic model of relevance to gastric physiology for generalization to other problems such as understanding the effects of acid in the upper gastrointestinal tract and the role of pH in the colonization of gut bacteria. Our results could further stimulate the development and testing of theoretical and computational fluid dynamics models to understand the motion of bacteria in gels and other confined geometries, as well as guide the design of artificial biomimetic swimmers using pH to control motility.

## MATERIALS AND METHODS

### *H. pylori* J99 culturing conditions

*H. pylori* strain J99 WT was provided by Prof. Sara Linden of University of Gothenburg, Sweden. For details of the strain and its original source see [41]. J99 was cultured initially from frozen stocks (stored in −80°C freezer) on Brucella agar (Brucella Medium Base, Oxoid, Basingstoke, Hampshire, England) supplemented with 10% sheep blood, 1% IsoVitox (Oxoid), 4 mg/L amphotericin B, 10 mg/L vancomycin and 5 mg/L trimethoprim for 48 hours then replated for an additional 48 hours on Brucella agar. Each culture was then inoculated in liquid media (BB10) containing 10% fetal bovine serum and 90% Brucella broth for 7 – 10 hours on the shaker, then diluted with broth and incubated for additional 12 – 16 hours on the shaker to optimize the number of motile bacteria. All agar or broth cultures were maintained at 37°C under microaerobic conditions in a tri-gas environment containing 5 – 12% CO_2_ and 5 - 15% O_2_ using BD GasPak systems (BD Biosciences, San Jose, CA, USA) consisting of a non-vented cylinder or a heavy Ziplock bag. The concentration of bacteria in liquid culture was monitored by measuring the absorbance using a spectrophotometer. The liquid culture is added to each sample when the bacteria reaches the exponential growth phase, when *OD*_*600*_ = 0.6 - 0.7.

### PGM preparation

PGM was collected from pig stomach epithelium and purified with Sepharose CL-2B column chromatography, followed by density gradient ultracentrifugation, described in Celli *et al* [7] and more fully in the original references therein. Lyophilized PGM was weighed, and the appropriate amount of PGM was dissolved in sterile water to prepare a final solution at 15 mg/ml after addition of 10% bacterial or bead sample and 10% buffer. Typically, 40 μl of sterile water was added to 0.75mg of PGM and mixed using a vortex mixer for 10 to 20 minutes, until there were no visible un-hydrated PGM (white spots). This solution was allowed to hydrate and equilibrate for 48 hours at 4°C and was used for a maximum period of one week. When ready to use, either 5 μl of 0.1 M phosphate-succinate buffer (adjusted to various pHs using HCl) or 5 μl of BB10 with HCl added to adjust pH and thoroughly mixed with a positive displacement pipette. The buffered PGM solution was incubated at 37°C for at least 40 minutes before 5 μl of bacterial or beads sample could be added.

### pH calibration in BB10 and PGM

BB10: For each pH, 100 μl of bacteria liquid culture or particle solution was added to 800 μl of fresh BB10, follow by a gradual addition of HCl and BB10 until a desired pH level and a final total volume of 1 ml were reached. A 3-point calibrated handheld pH meter was used to monitor the pH of each sample.

PGM: For each pH, 2.5 μl of bacteria liquid or particle solution was added to 20 μl of PGM solution in buffer, followed by a gradual addition of buffer and HCl and BB10 until a desired pH level and a final total volume of 25 μl. The pH of each sample was monitored using Hydrion pH strips. We noted over pH 2-6 the final pH of the PGM solution was higher than the initial buffer pH: at external buffer pH of 2, 4, 5, 6, 7 the PGM solution had pH 3.7, 4.6, 5.5, 6.1 and 6.7, respectively [27].

### Microrheology: sample preparation, fluorescence particle imaging, particle tracking

#### Sample preparation

Fluorescent particles of 1.001 μm diameter (Polysciences) were added to each sample to obtain 0.054% of final particle concentration by volume for all microrheology measurements. Samples were incubated at 37°C for 5-10 minutes prior to each measurement. After the incubation, 10 μl volume of each solution was pipetted onto a standard glass microscope slide with a spacer (Secure Seal Imaging Spacer, 9 mm in diameter, 0.12 mm in depth) and sealed with a coverslip.

#### Fluorescence imaging

The movies were acquired immediately at room temperature (∼21°C) with an Olympus I70 inverted optical microscope equipped with a 40x phase contrast lens (0.65 NA) and an Andor Zyla 5.5 sCMOS camera at 100 fps and 6.5 μm per pixel size. Fluorescent particles were excited using Olympus BH2 Mercury arc source. Imaging focus was optimized at the center Z positions of each sample to minimize edge effects. Three-second videos were obtained from several different x-, y-positions with optimal z-position throughout each sample. *Particle tracking:* The Brownian motion of the particles was tracked in MATLAB [v7.12.0] using PolyParticleTracker routine which locates the center of mass of each tracked object with a polynomial fit of the photon intensity around the object and weighted by a Gaussian function [42]. Mean square distance (*MSD*), viscosity (*η*), diffusion exponent (*α*), heterogeneity (*HR*) of each sample were calculated based on the tracker position output. The number of trajectories analyzed are given in Fig. S1. As seen in Fig. S1, the MSD for individual trajectories eventually exhibits a decline as particles move out of the image plane and the out of plane displacement cannot be measured. We used this point as a cut off in analyzing particle trajectories.

### Motility: phase contrast microscopy live cell imaging, cell tracking

#### Sample preparation

Each PGM or BB10 sample was incubated at 37°C for 45 minutes. Bacteria were cultured in liquid broth (BB10) to an *OD*_*600*_ of 0.6-0.7 then added to each sample to produce a 10% bacteria mixture by volume. The bacteria mixture was incubated at 37°C under microaerobic conditions using GasPak systems and constant agitation for 45 minutes before measurement. A 10-μl volume of each sample was applied to a glass microscope slide with a secure seal spacer and sealed with a coverslip.

#### Phase contrast imaging

The samples were imaged immediately at room temperature using an Olympus IX70 inverted phase contrast microscope equipped with a 40x objective lens (0.65 NA), a halogen light source, and an Andor Zyla 5.5 sCMOS camera at 33 fps and 6.5 μm pixel size. Videos of bacteria swimming in mid-plane between the coverslip and microscope glass slide were acquired for 9 s using Micro-Manager open source acquisition software. All imaging experiments at different pH in the two media, BB10 and PGM were conducted from the same batch of bacteria to minimize variation from batch to batch. However, this sequential approach limited the number of different pHs that could be examined as bacteria cannot be used for very long. To image body rotation we used 100x oil-immersion objective lens (1.25 NA) and an Andor Zyla 5.5 sCMOS camera at 100 or 200 fps. Three-second videos of bacteria swimming in mid-plane between the coverslip and microscope glass slide were acquired.

#### Cell tracking and analysis of trajectories

Bacteria trajectories were tracked using PolyParticleTracker MATLAB routine to determine the instantaneous position in 2 dimensions. Using the position vector of each bacteria we calculate the instantaneous speed as the displacement per unit time between two consecutive frames and the direction in which the bacterium is traveling as the angle φ(*t*) of the 2-dimensional velocity vector. Further analysis was done by segmenting each trajectory into runs and reorientations using a modified version from Hardcastle [37] of the method developed by Theves *et al* [43].

Reorientation events were determined by looking for large changes in the maximum of |Δ*ϕ*| and/or the minimum of *v* according to:

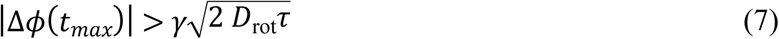

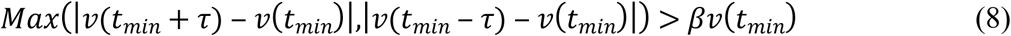

where *t*_*max*_ is the time at which |Δ*ϕ*| is maximum; *t*_*min*_ is the time at which *v* is minimum; *τ* = 0.03 s is the time between frames; *γ* is a threshold variable that determines how much larger than the angular deviation 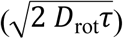 that |Δ*ϕ* (*t*_*max*_)| has to be; and *β* is a threshold variable that determines how much larger than *v*(*t*_*min*_) that the speed change has to be to be considered a reorientation event. We estimated the rotational diffusion constant *D*_*rot*_ of a bacteria by approximating it is an ellipsoid with semi-minor axis *a* ≈ 0.5 μm and semi-major axis *b* ≈ 1.5 μm,

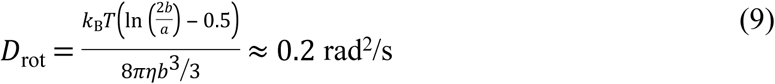

Using this rotational diffusion constant and by locating *v*(*t*_*min*_), we found γ=8.5 and *β* = 1.75 to be sufficient to identify reorientations events where bacteria actively reoriented their swimming direction.

The bacterium was assumed to stay in the reorientation state for a time *tre* calculated by examining if the local angle changes |Δφ(*t*\s\do5(*max*) + *t*\s\do5(*re*))| was large compared to the |Δφ(*t*\s\do5(*max*))| and if the displacement of the bacterium was smaller than that of Brownian motion, using the criterion:

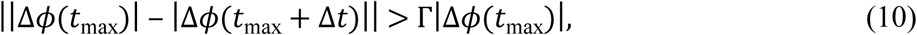

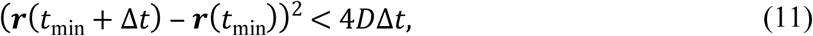

where 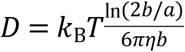 is the translational diffusion of an ellipsoid doing Brownian motion and Γ is the percentage change in angle for which the bacterium is still considered to be in reorientation event. We found that Γ = 0.7 made the best identification. For *H. pylori D =* 0.26 μm^2^/s. Reorientation angles between runs are denoted by θ_re_ and if larger than 140° they were identified as reversals. The reversal frequency was obtained by ensemble averaging of the number of reversals divided by the time duration of each trajectory. Reversal percent was defined as the ratio of the number of angle changes greater than 140° to the total number of angle changes.

### Body rotation and cell shape analysis

The movie of each bacterium was individually and manually cropped using ImageJ. The cell body contour and the centerline of each bacterium were extracted and aligned using CellTool python software [33]. For each trajectory we can obtain numerous images of the cell shape of an individual bacterium and we select the one with the largest axial length (most in-plane image) as explained in Constantino *et al* [31] to obtain the cell shape parameters. The body rotation rate of each bacterium was measured by monitoring the change in the alignment angle of the bacterium and the time between two maximum points, as described in Constantino *et al* [31].

### Numerical analysis of distributions and model calculations

All numerical analysis was done using Matlab.

### Two peak fitting analysis of speed distributions

The speed distribution curves for both *v*_*ins*_ and *v*_*run*_ were fit to a sum of two Gaussians using

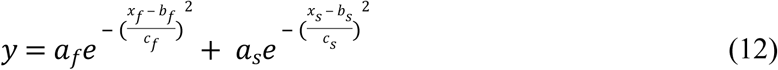

where *a, b* and *c* denote the amplitude, position and width of the fast (*f*) and slow (*s*) peaks, and *x* denotes the variable of interest, *v*_*ins*_ or *v*_*run*_. Note that the width *c* is equal to *√2.σ*, where *σ* is the standard deviation of the Gaussian distribution.

### Resistive Force Theory calculations for estimation of torque

For swimming bacteria, motor torque *T*_*m*_ was estimated from shape factor *S*_*h*_ and bacterium swimming speed *v*_*h*_ for each bacterium imaged at 100X in PGM and BB10 at different pH.

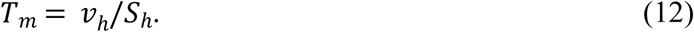

The shape factor *S*_*h*_ was calculated using helical cell body and helical flagella bundle drag coefficients [31]

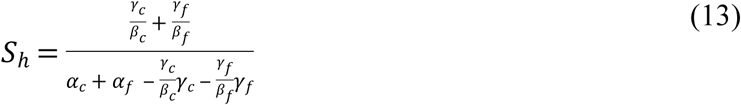

where *c* denotes helical cell body with helical pitch *λ*_*c*_, radius of helical filament *a*_*c*_, helical radius *R*_*c*_, pitch angle Φ_*c*_ given by *tan* Φ_*c*_ = 2*πR*_*c*_/ *λ*_*c*_, helical contour length *L*_*c*_ = *X*_*L*_/cos Φ_*c*_, *X*_*L*_ is the axial length of cell body. The hydrodynamic translational, rotational and propulsion drag coefficients for the cell body, related to the local normal and tangential force *C*_*n*_, *C*_*t*_ are given below:

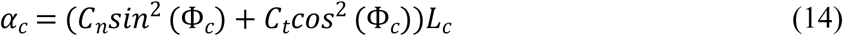

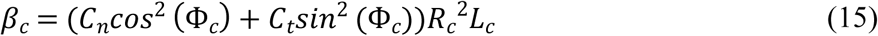

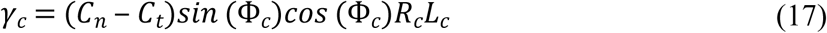

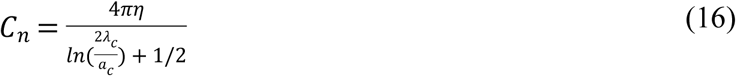

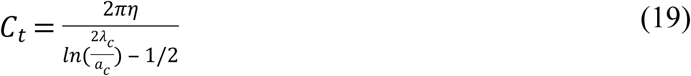

Here *η* denotes the viscosity of the medium. By changing the subscript *c* to *f* these equations give the same quantities for the flagella helix in terms of its helical parameters. The flagella parameters that we used were same as Constantino *et al* [31] : *X*_*f*_ = 2.97 μm, *a*_*f*_ = 0.07μm, *λ*_*f*_ = 1.58μm and *R*_*f*_ = 0.14μm.

The cell body torque can be calculated from the measured swimming speed

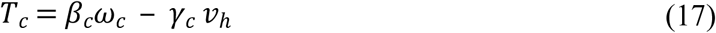

For stuck bacteria which rotate but do not translate *v*_*h*_ is zero and simplifies to

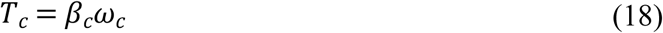

## ACKNOWLEDGEMENTS

We thank Prof. Henry Fu (Univ. of Utah) and Prof. Maria Kamenetska (Boston University) for helpful discussions.

## SUPPORTING INFORMATION

**Figure S1A**. All particle MSD vs delay time in BB10 at different pHs as indicated. The average <MSD> and relative error of log MSD defined as ε = ±0.436 σ_MSD_/<MSD> are displayed by the black line and error bars. Numbers in parenthesis on pH legends indicate number of particles tracked. Dashed lines of slope 1 and slope 0.5 are indicated to guide the eye.

**Figure S1B**. All particle MSD vs delay time in PGM at different pHs as indicated. The average <MSD> and relative error of log MSD defined as ε = ±0.436 σ_MSD_/<MSD> are displayed by the black line and error bars. Numbers in parenthesis on pH legends indicate number of particles tracked. Dashed lines of slope 1 and slope 0.5 are indicated to guide the eye.

**Figure S2**. Two peak fit of distribution of instantaneous speeds, *v*_*inst*_ for J99 *H. pylori* bacteria swimming in BB10 and PGM at different pHs as indicated. The number of motile trajectories that were analyzed ranges from 185 at pH 4 to 388 at pH 3 in BB10 and from 136 at pH 4 to 625 at pH 4.5 in PGM.

**Movies** (see separate files for movies)

**Movie S1.** Phase contrast microscopic video of a bacterium at 100X swimming in BB10 at pH 4, showing forward and reverse motions, as described in the text.

**Movie S2.** Phase contrast microscopic video of a bacterium rotating about a fixed point in PGM gel at pH 4 with slow and fast body rotation rates over time.

**Movie S3.** Phase contrast microscopic video of a bacterium translating in a fixed circular trajectory in PGM at pH 4.

**Movie S4.** Phase contrast microscopic video of a bacterium in PGM at pH 4.5 showing rotations with random translational motions.

**Movie S5.** Phase contrast microscopic video of a bacterium in BB10 at pH 4 showing translational displacement while rotating.

